# A rapid visuomotor response on the human upper limb is selectively influenced by implicit, but not explicit, motor learning

**DOI:** 10.1101/354381

**Authors:** Chao Gu(顾超), J. Andrew Pruszynski, Paul L. Gribble, Brian D. Corneil

## Abstract

How do humans learn to adapt their motor actions to achieve task success? Recent behavioral and patient studies have challenged the classic notion that motor learning arises solely from the errors produced during a task, suggesting instead that explicit cognitive strategies can act in concert with the implicit, error“based, motor learning component. Here, we show that the earliest wave of directionally-tuned neuromuscular activity that occurs within ~100 ms of peripheral visual stimulus onset is selectively influenced by the implicit component of motor learning. In contrast, the voluntary neuromuscular activity associated with reach initiation, which evolves ~100 to 200 ms later is influenced by both the implicit and explicit components of motor learning. The selective influence of the implicit, but not explicit, component of motor learning on the earliest cascade of neuromuscular activity supports the notion that these components of motor learning can differentially influence descending motor pathways.

## INTRODUCTION

Motor learning occurs throughout the human lifespan, from children learning to walk to the aged adjusting to a new set of reading glasses. Motor learning involves establishing and constantly recalibrating the mapping of a desired goal onto the required motor commands [1]. A predominate theory of motor learning posits that learning arises from an *implicit* error-based process, in which the brain learns by computing an error between actual and predicted sensory consequences of the generated motor command [2,3]. Recent behavioral work using a visuomotor rotation task [4] which systematically rotates the visual cursor denoting hand position around the center of the workspace, has suggested that a second *explicit* process also contributes to motor learning [5–7]. The explicit process is driven by awareness of task errors, which participants exploit to achieve task success. Research with individuals who have brain lesions shows that the implicit and explicit components of motor learning have distinctive neural substrates, relying on the integrity of cerebellar [8,9] and frontal circuits [10,11], respectively. However, multiple descending pathways originating from the cortex and brainstem contribute to motor control in healthy individuals [12–14] and the comparative influence of the implicit and explicit components of motor learning on these pathways is not known.

Our interest here is to compare the effect of motor learning on the first wave of directionally-tuned upper limb muscle activity that occurs time-locked ~100 ms after visual stimulus onset (termed *stimulus-locked responses*, or *SLRs*) [15] to the muscle activity that occurs at the time of reach initiation, roughly ~200-300 ms after stimulus onset [16]. Previous work has shown that the largest SLRs occur when stimuli are presented at locations associated with the largest reach-related responses [15,17], and SLRs persist even if the reach movement is withheld [18,19] or proceeds in the opposite direction [20]. These response properties, as well as the fact that SLRs evolve at latencies that preclude extensive cortical processing, have led us to propose that SLRs and later reach-related activity arise from distinct descending motor pathways [15,20].

Here, we study how the implicit and explicit components of motor learning influence these two waves of EMG activity during the visuomotor rotation task. Success in this task requires that participants learn a new mapping between the location of the visual stimulus and the direction of the reach movement. We quantify the change in directional tuning of the SLR and reach-related activity across three different variants of the visuomotor rotation task that either combine or isolate the implicit and explicit components of motor learning. We show that changes in SLR tuning only occur during tasks that involve implicit motor learning, and that the partial shifts in SLR tuning observed during these experiments (~10-15° for rotations of both 40 and 60°) are consistent with previous estimates of implicit learning based on verbal reports of participants’ explicit aiming direction [6,21]. In contrast, the tuning of reach-related activity shifts completely in all tasks, consistent with influences of both implicit and explicit motor learning. Taken together, our results show that the earliest wave of muscle activity following a visual stimulus is selectively influenced by implicit motor learning, whereas later voluntary waves of muscle activity are influenced by both implicit and explicit motor learning.

## RESULTS

In all three experiments, participants (*N* = 8, 14, and 18, respectively, 4 participants performed in multiple experiments) sat at a desk and used their right hand to interact with the handle of a robotic manipulandum that controlled the position of a cursor, presented on a horizontal mirror reflecting a downward facing LCD screen (**METHODS**). The participant’s right arm was occluded throughout all experiments; thus, the position of the cursor was the only visual cue of the manipulandum presented to the participants. The visuomotor rotations in Experiments 1 and 2 were introduced by rotating the visual feedback of the cursor around the central starting position (**Fig. 1d**). In all three experiments, we measured both the *x*- and *y*-positions of the manipulandum and the EMG activity from the right pectoralis major (PEC) muscle while participants performed right-handed reach movements to one of eight peripheral stimuli equally spaced 10 cm around the starting position.

**Figure 1:**
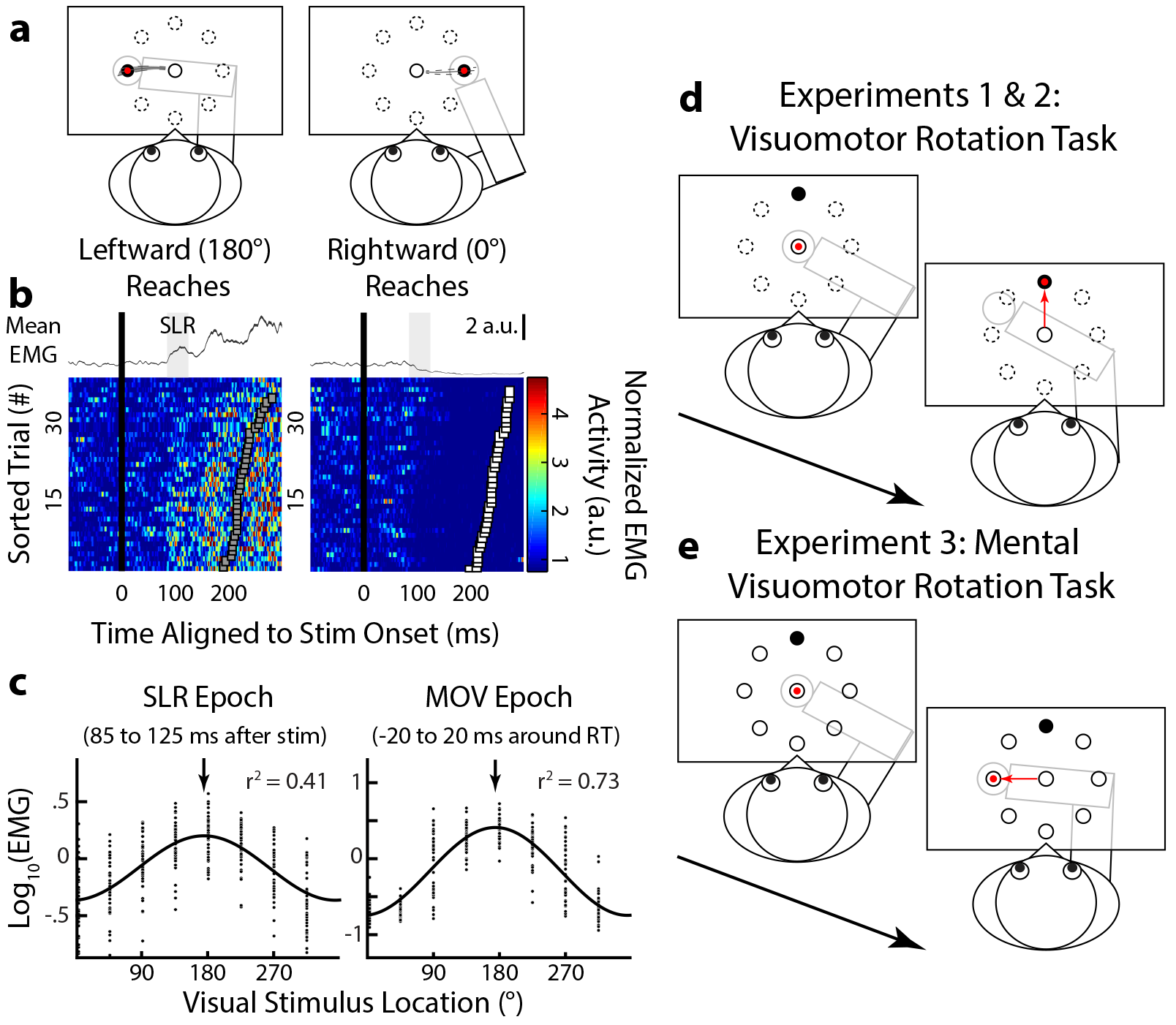
Experimental paradigm and spatial tuning of the stimulus-locked response (SLR) on human limb muscle during visually-guided reaches. **a.** The mean ± SD normalized movement trajectories for leftward and rightward visually-guided reach for a representative participant. **b.** The corresponding mean ± SEM (top panels) and individual trials (bottom) of EMG activity from the right pectoralis major muscle aligned to visual stimulus onset (black line). For the color panels, each row represents EMG activity from a single trial, with trials sorted based on reach RT (squares). EMG activity diverged during the SLR epoch (shaded regions, 85-125 ms after stimulus onset), regardless of the ensuing RT. **c.** Sinusoidal relationship between the normalized mean EMG activity and visual stimulus location during the SLR (left panel) and MOV (right) epochs for this participant. Arrows indicate the PD of each fit. **d.** Experiments 1 and 2: the visuomotor rotation task. Participants generating reach movements to move the cursor (red circle) to the visual stimulus location (black circle). To induce motor learning, the cursor was systematically rotated (60° CW in this case) around the start position. **e.** Experiment 3: the mental rotation task. During the task, the cursor always gave veridical feedback of the robotic handle but participants were explicitly instructed to reach to the stimulus location 90° CCW to the visual stimulus location.

**Figure 1a** shows the normalized mean ± SD movement trajectories for both the leftward (180° CCW from straight right) and rightward (0°) stimulus locations from a representative participant, when they had veridical visual feedback of their hand position (i.e., the cursor moved in register with the participant’s hand). **Figure 1b** shows the corresponding normalized mean ± SEM (top) and individual (bottom color panels) PEC EMG activity from leftward and rightward trials. EMG activity was aligned to the onset of the peripheral visual stimulus onset (thick black vertical lines), and individual trials were sorted based on reaction time (RT; squares, fastest to slowest from bottom to top). We observed a reliable SLR, which consisted of a brief increase or decrease in EMG activity ~100 ms after the presentation of leftward or rightward stimulus locations, respectively [15,18,20]. We defined the SLR magnitude for each trial as the mean EMG activity during the SLR epoch (85-125 ms after stimulus onset, shaded regions in mean EMG sub-panels in **Fig. 1b**).

To determine the directional tuning of the EMG activity during both the SLR and the later reach-related response (MOV, −20 to 20 ms around RT) epochs, we derived the preferred direction (PD) of each epoch assuming a sinusoidal fit (**Eq. 1**). **Figure 1c** shows the log-normalized EMG activity as a function of visual stimulus location (arrows indicate the PDs of each fit). With veridical feedback, a reliable SLR was detected in 29 out of 32 participants (see *ROC analysis* in **METHODS** for detection criteria). Consistent with a previous study [15], we also found a small but reliable difference in PD of EMG activity between the SLR and MOV epochs (mean ± SEM: 172.5° ± 1.6° and 180.0° ± 1.2°, respectively, paired *t*-test, *t*_36_ = −4.0, *P* = 0.001). Data from participants who did not exhibit an SLR were excluded from all subsequent analyses (see **METHODS** for exact numbers for each experiment). Having established the tuning of EMG activity during the SLR and MOV epochs with veridical hand position feedback, we next examined how the PDs changed during two different visuomotor rotation tasks (**Fig. 1d**) and a mental visuomotor rotation task (**Fig. 1e**).

### Partial adaptation of the SLR during an abrupt 60° CW visuomotor rotation

In Experiment 1, we used an abrupt visuomotor rotation task which has been previously shown to engage both implicit and explicit motor learning components [5,6]. During both the Pre- and Post-Rotation blocks (**Fig. 2a**, black and blue shades, respectively), participants (*N* = 7) performed 60 and 80 cycles (a cycle consists of 8 reaches, 1 reach per direction) of visually-guided reaches under veridical visual feedback, respectively. During the Peri-Rotation block (red, 80 cycles), we imposed a 60° CW rotation on the visual cursor around the start position. **Figure 2a** also shows the group mean ± SEM reach endpoint (white dot and shade) plotted relative to the stimulus location, while the solid black line indicates perfect task performance. Consistent with previous experiments [22,23], our participants rapidly adapted their endpoint reach direction during the beginning of the Peri-Rotation block and exhibited signs of implicit learning as seen by the aftereffect during the beginning of the Post-Rotation block [5]. We excluded the first 20 cycles of both the Peri- and Post-Rotation blocks to ensure that participants’ behavioral performance had plateaued. We observed an increase in median RTs during the Peri-Rotation block (**Fig. 5a**, group mean ± SEM = 301 ± 17 ms) compared to either blocks with veridical feedback (Pre- and Post-Rotation, 246 ms ± 14 ms and 254 ± 13 ms, paired *t*-test, *t*_6_ = −7.5 and −3.4, *P* = 0.001 and *P* = 0.01, respectively). Prolonged RTs during the visuomotor rotation task have been associated with explicit motor learning as participants employ an aiming strategy [24,25]. Thus, participants’ behavior provided evidence for the engagement of both implicit and explicit motor learning components during this task.

**Figure 2:**
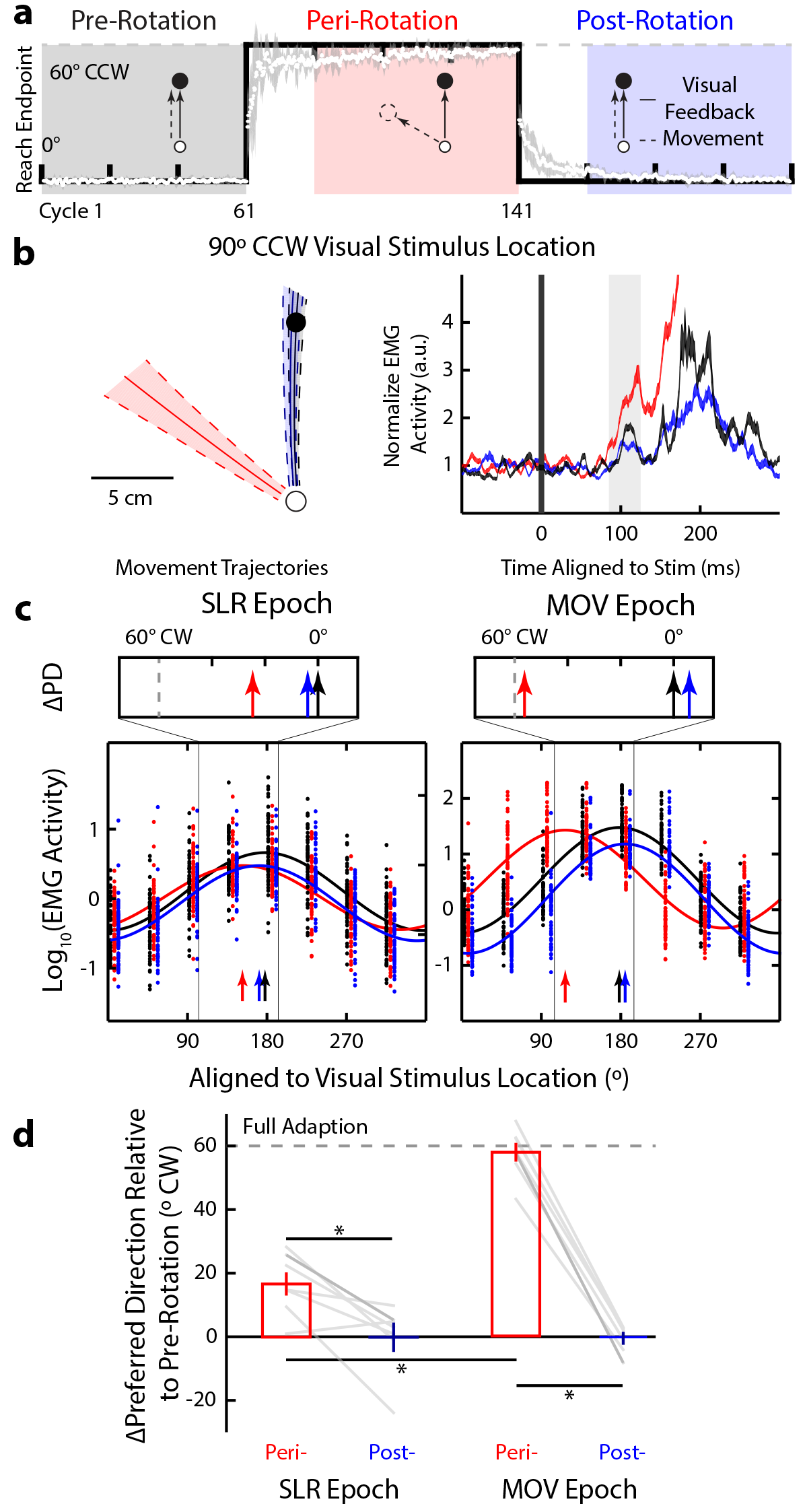
Partial adaptation of the SLR tuning during the abrupt visuomotor rotation task. **a.** Timeline and behavioral performance during an 60° CW abrupt visuomotor rotation. The group mean ± SEM (white circles and gray shade) reach endpoint per cycle relative to the stimulus location is plotted against perfect task performance (black line). Veridical visual feedback was provided during Pre- (black shade) and Post-Rotation (blue) blocks. During the Peri-Rotation (red) block, the virtual cursor feedback was rotated around the start position by 60° CW. **b.** Mean ± SD normalized movement trajectories and mean ± SEM PEC EMG activity for the outward visual stimulus location (90° CCW from straight right) of a representative participant. The EMG activity is aligned to stimulus onset, and the SLR epoch (85-125 ms after stimulus onset) is highlighted. **c.** Sinusoidal tuning curve fits (**Eq. 1**) between visual stimulus location and the normalized mean EMG activity during the SLR (left panel) and MOV epochs (right). Each dot indicates data from a single trial, while the solid lines shows the best fit for each block; vertical arrows indicate the PDs for each fit. Note for illustration purposes only, we have staggered the individual trial data. Top inserts show the shifts in PD (*ΔPD*) during the Peri- and Post-Rotation blocks relative to the Pre-Rotation block. Vertical dashed gray line represents full adaptation to the 60° CW visuomotor rotation. **d.** Group mean ± SEM of *ΔPD* for both Peri- (red bars) and Post-Rotation blocks (blue) during both the SLR and MOV across all participants. A *ΔPD* = 0° or *ΔPD* = 60° CW would indicate either no adaptation or a complete adaptation to the imposed rotation, respectively. Each gray line represents data from an individual participant, with the darker line indicating data from the participant in **c.** \**P* < 0.05.

**Figure 2b** shows mean movement trajectories and PEC EMG activity for the outward visual stimulus location (90° CCW) across the three different blocks, for one participant. As seen from the mean movement trajectories, during Peri-Rotation (red) the participant learned that the imposed 60° CW visuomotor rotation required them to generate a left-outward reach movement ~60° CCW to the stimulus location. These left-outward movements during the Peri-Rotation block required more PEC recruitment compared to straight outward movements during both Pre- and Post-Rotation blocks. As expected, during the MOV epoch we observed reliable modulation in PEC EMG activity across blocks (1-way ANOVA, main effect, *F*_(2,176)_ = 486.4, *P* < 10^−71^), with greater EMG activity during Peri-compared to both Pre- and Post-Rotation (post-hoc Tukey’s HSD, both *P* < 10^−9^).

We also observed a similar pattern of modulation during the SLR epoch (1-way ANOVA, main effect, *F*_(2,176)_ = 7.97, *P* = 0.001), with greater EMG activity during the SLR epoch for Peri-compared to both Pre- and Post-Rotation blocks (post-hoc Tukey’s HSD, *P* = 0.006 and *P* = 0.001, respectively). Thus, even though the same visual stimulus location was presented across all three blocks, the magnitude of the SLR changed during motor learning.

To quantify the influence of motor learning on directional tuning, we derived the PDs of EMG activity during the two different epochs for all three blocks (colored arrows in **Fig. 2c**). We normalized the results across participants by using each participant’s PD during the Pre-Rotation block as a baseline and quantified the shifts in PD (ΔPD) for both Peri- and Post-Rotation blocks (top panels in **Fig. 2c**). Across participants (**Fig. 2d**), we found that ΔPD for the MOV epoch adapted almost completely during the Peri-Rotation block (ΔPD mean ± SEM = 57.7 ± 2.9° CW, one sample *t*-test, *t*_6_ = 19.61, *P* < 10^−5^) to the imposed 60° CW visuomotor rotation (gray dashed line). Note this is expected as we aligned the tuning curves relative to visual stimulus location rather than the reach direction. We also found that ΔPD returned to baseline during the PostRotation bock (ΔPD = 0.7 ± 1.6° CW, one sample *t*-test, *t*_6_ = 0.46, *P* = 0.66), and a reliable difference in ΔPD between the Peri- and Post-Rotation blocks (2-way ANOVA ‒ epoch and rotation blocks, interaction effect, *F*_(1,24)_ = 41.63, *P* < 10^−6^, post-hoc Tukey’s HSD, *P* < 10^−8^). Thus, we observed nearly complete adaptation (ΔPD ≈ 60° CW) and de-adaptation (ΔPD ≈ 0° CW) during the MOV epoch for the Peri- and Post-Rotation blocks, respectively.

We next examined the change in the directional tuning of EMG activity during the SLR epoch. Like the later MOV epoch, we also observed reliable adaptation during the Peri-Rotation block (ΔPD = 16.7 ± 3.6° CW, one-sample *t*-test, *t*_6_ = 4.6, *P* = 0.004), and de-adaptation during the Post-Rotation block (ΔPD = 0.0 ± 4.2° CW, one-sample *t*-test, *t*_6_ = 0.01, *P* = 0.99). However, the extent of adaptation during Peri-Rotation for the SLR epoch was reliably smaller than that during the later MOV epoch (2-way ANOVA ‒ epoch and rotation blocks, post-hoc Tukey’s HSD, Peri-Rotation ‒ SLR vs MOV epoch, *P* < 10^−7^).

To summarize the results from Experiment 1, motor learning induced via an abrupt 60° CW visuomotor rotation systematically altered the tuning of the SLR, despite its short-latency. However, unlike the full adaptation of EMG in the later MOV epoch, we observed only partial adaptation of EMG during the SLR interval. The abrupt visuomotor rotation task is thought to engage both implicit and explicit motor learning components. In Experiment 2 we tested whether the shift in SLR tuning is still present when the explicit component of motor learning is minimized or eliminated.

### SLR adaptation occurs despite a lack of explicit awareness of a visuomotor rotation

In Experiment 2, participants (*N* = 14) performed a gradual visuomotor rotation task (**Fig. 3a**). A previous imaging study has suggested that abrupt and gradual visuomotor rotation tasks engage different neural substrates [26], and behavioral studies have shown that gradual visuomotor rotations produced larger aftereffects [27] and longer-lasting retention [28] compared to abrupt visuomotor rotations. In Experiment 2, we imposed a visuomotor rotation gradually (1° per cycle). Once again, participants initially performed visually-guided reaches to one of eight equidistant visual stimuli with veridical feedback (**Fig. 3a**, Test Block 1, Pre-Rotation) for 40 cycles. Then for the next 20 cycles, the visual feedback of the cursor was rotated either 1° CW or CCW per cycle (solid or dashed lines), counterbalanced between participants. Over the next 40 cycles, the visual feedback remained rotated at 20° CW or CCW (Test Block 2). Afterwards, the feedback was rotated 1° per cycle in the opposite direction to the initial imposed rotation for 40 cycles. Finally, the feedback remained constantly rotated at 20° CCW or CW (Test Block 3). We found no reliable differences in endpoint reach direction between the three Test Blocks based on the order of imposed rotation (2-way ANOVA, Test Blocks and group, main effect of group, *F*_(2,36)_ = 0.07, *P* = 0.93). Thus, we pooled data from all participants together for the subsequent analyses.

**Figure 3:**
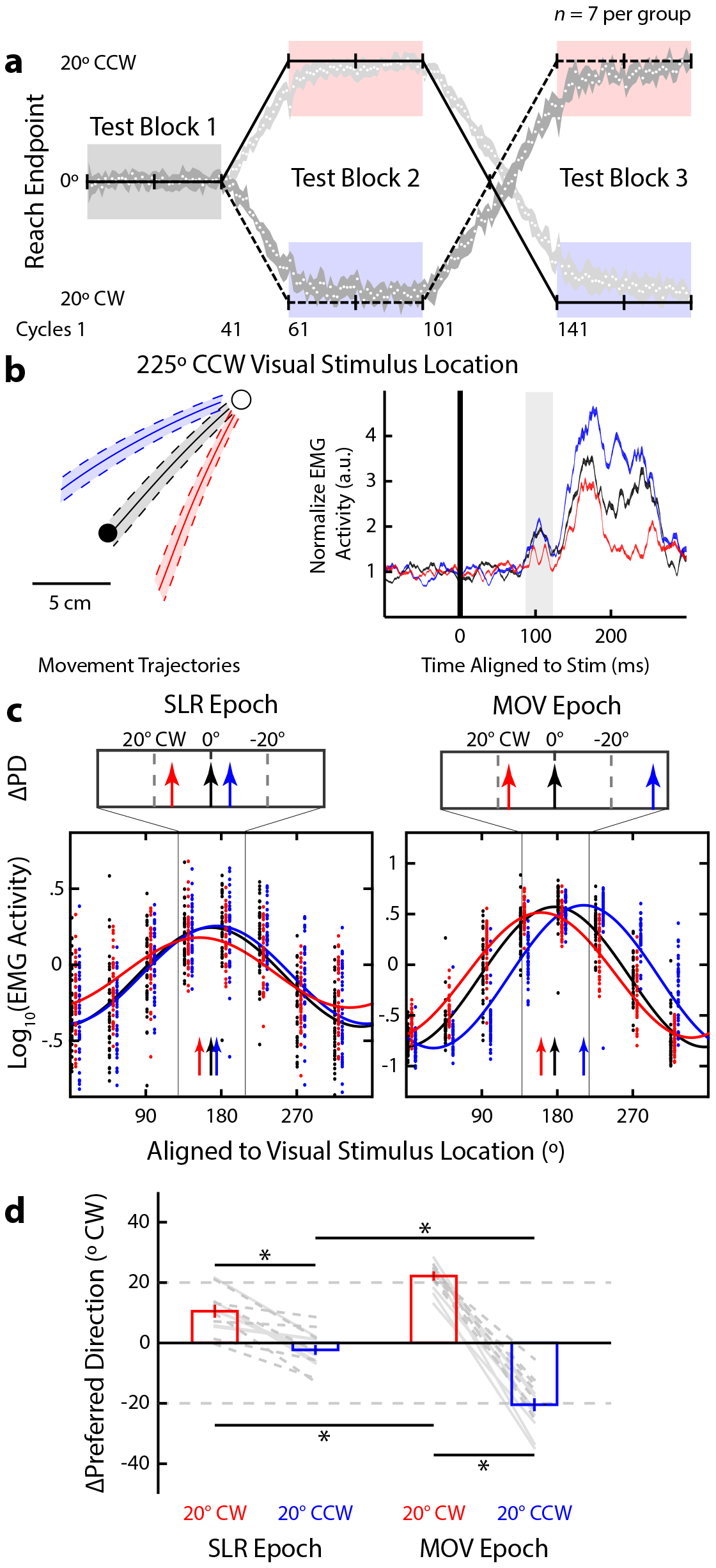
Partial adaptation of the SLR tuning during the gradual visuomotor rotation task. Same layout as **Fig. 2. a.** Timeline and behavioral performance during a gradual visuomotor rotation task. After the 40 cycles of reaches (Test Block 1) with veridical cursor feedback, the cursor was gradually rotated 1° per cycle to 20° CW (black solid line) or CCW (dashed line). After participants performed 40 cycles with the cursor constantly rotated 20° CW or CCW (Test Block 2), the cursor was rotated in the opposite direction for 40 cycles. Finally, participants performed 40 cycles with the cursor constantly rotated 20° CCW or CW (Test Block 3). Both groups performed reaches with veridical (Pre-Rotation, black), 20° CW (red), and 20° CCW (blue) visual feedback blocks. **b.** Mean ± SD movement trajectories and mean ± SEM EMG activities for the left-inward visual stimulus location (225° CCW) during the three blocks from a participant who experienced the CW rotation first. **c.** PD for each of the Test Blocks during both the SLR and MOV epochs (vertical arrows). **d.** Mean ± SEM of the ΔPD for CW and CCW blocks compared to Pre-Rotation block for both the SLR and MOV epochs across all participants. Dashed or solid lines indicate participants who first experienced CW or CCW rotation, respectively. * *P* < 0.05.

The size of the imposed visuomotor rotation, 1° per cycle, during Experiment 2 is less than the trial-by-trial variance of the participants’ reach endpoint during the Pre-Rotation block (Gaussian fit, mean ± SD, μ = 0.4 ± 0.1, *σ*^2^ = 5.0 ± 0.2, adjusted r^2^ = 0.94 ± 0.01). Consistent with previous studies [29,30], participants reported no explicit awareness of changes in the underlying sensorimotor mapping at any point during the experiment. Further, unlike Experiment 1, we found no difference in median RTs between veridical feedback (**Fig. 5b**, Pre-Rotation, mean ± SD = 232 ± 5 ms) and the two rotation blocks (CW and CCW, 233 ± 5 ms and 236 ± 5 ms, paired *t*-test, *t*_13_ = −0.65 and −1.48, *P* = 0.52 and *P* = 0.16, respectively). This lack of RT increase during the gradual visuomotor rotation is also consistent with a minimal influence of explicit aiming during the experiment.

**Figure 3b** shows mean movement trajectories and PEC EMG activity for one participant, for the left-inward stimulus location (225° CCW) across the three Test Blocks: Pre-Rotation, 20° CW, and 20° CCW (black, red, and blue traces, respectively). Like in Experiment 1, we found reliable differences in normalized EMG activity across the three blocks for both the SLR and MOV epochs for this stimulus location (1-way ANOVA, main effect, *F*_(2,109)_ = 5.74 and 57.6, *P* = 0.004 and *P* < 10^−17^, respectively). For example, during the 20° CW rotation block, the participant generated reaches away from the PD of the PEC muscle, hence there was a decrease in mean EMG activity both during the MOV epoch (red trace in **Fig. 3b**, starting after ~150 ms after stimulus onset post-hoc Tukey’s HSD, *P* < 10^−5^) and during the SLR epoch (shaded region, post-hoc Tukey’s HSD, *P* = 0.01). **Figure 3c** shows the tuning curve fits during both the SLR and MOV epochs across the three different blocks for this participant, demonstrating the changes in the PD in both the SLR and MOV epochs for this participant.

When we examined the shifts in PD across our sample, as expected we observed full ΔPD adaptations of 22.2 ± 1.1° CW and 20.4 ± 2.1° CCW during the MOV epoch for the 20° CW and 20° CCW rotation blocks relative to the Pre-Rotation block, respectively (**Fig. 3d**, right panel, 2-way ANOVA ‒ Epoch and Rotation, interaction effect, F_(1,52)_ = 77.9, *P* < 10^−11^, post-hoc Tukey’s HSD, *P* < 10^−8^). When we performed the same analysis during the SLR epoch (**Fig. 3d**, left panel), we found that the SLR ΔPD rotated 10.5 ± 1.7° CW and 2.3 ± 1.6° CCW for the 20° CW and CCW rotation, respectively (post-hoc Tukey’s HSD, *P* < 10^−4^). As in Experiment 1, we observed a reliable smaller overall change in ΔPD during the SLR versus MOV epoch when collapsing these changes across the 20° CW and 20° CCW rotation blocks (12.8 ± 1.9° and 42.6 ± 2.1°, paired *t*-test, *t*_13_ = 11.0, *P* < 10^−7^).

Thus, as with an abrupt visuomotor rotation, motor learning induced by a gradual visuomotor rotation systematically altered the tuning of the SLR. Experiment 2 also demonstrated that explicit awareness of changes in the underlying visuomotor mapping is not required for the tuning of the SLR to change. However, the extent of adaptation during the SLR epoch was still reliably less than that observed in the later MOV epoch. This finding is consistent with literature suggesting that a cognitive strategy may still be engaged in the gradual visuomotor rotation task, despite the lack of explicit awareness [30].

### Changes in the explicit aiming strategy do not alter the PD of the SLR

In Experiment 3 participants (*N* = 13) performed a mental visuomotor rotation task [5,31]. Unlike in the first two experiments, participants received veridical visual feedback of their hand position throughout the experiment. It has been proposed that this eliminates implicit motor learning, since such learning is thought to occur only when there is a mismatch between the visual location of the virtual cursor and the participant’s hand position [5,9]. Instead, participants were explicitly instructed to reach either directly to the stimulus location (VIS block, **Fig. 4a**, grey) or 90° CCW relative to the stimulus location (Rotation [ROT] block, red). The order of the blocks was counterbalanced between participants. To assist participants, all eight stimulus locations were presented as open circles throughout the whole experiment, and the peripheral stimulus onset occurred when one of the open circles filled in. Like in Experiment 1, we found an increase in median RTs during the ROT (**Fig. 5c**, mean ± SEM = 398 ± 15 ms) compared to VIS Block (243 ± 7 ms, paired *t*-test, *t*_12_ = −17.8, *P* < 10^−9^), supporting the idea that participants used an aiming strategy during the ROT block.

**Figure 4:**
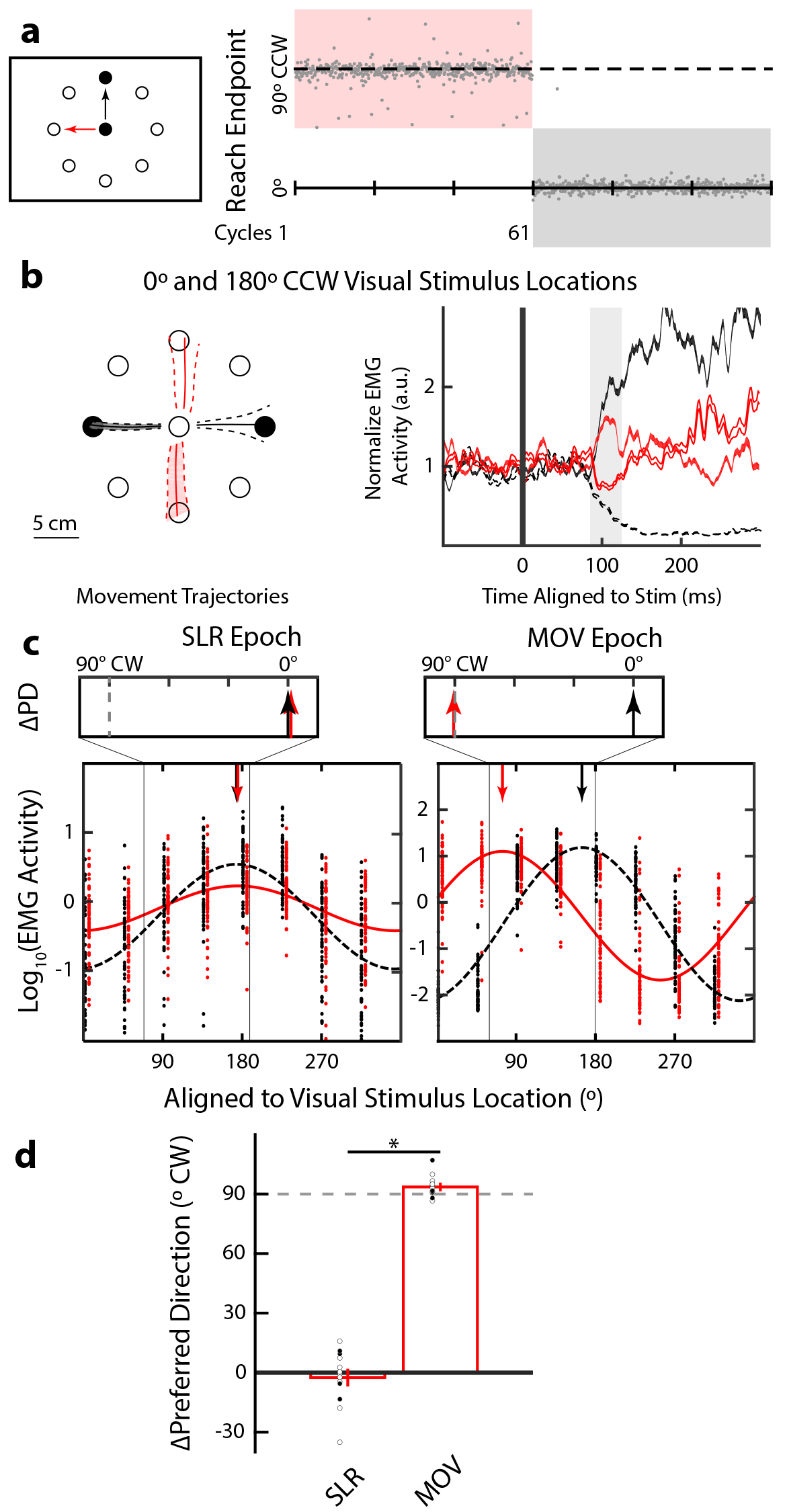
SLR tuning did not adapt during a mental visuomotor rotation task. Same layout as Fig. 2. a. Task schematic, timeline and behavioral performance for a representative participant during the mental visuomotor rotation task. Veridical visual feedback was given throughout the whole experiment. Participants were instructed to reach directly (VIS, black) or 90° CCW (ROT, red) to the stimulus location, with the order was counterbalanced across participants. **b.** Mean ± SD movement trajectory and mean ± SEM EMG activity for both the leftward and rightward stimulus locations. **c.** PD for each both the VIS and ROT blocks during both the SLR and MOV epochs (vertical arrows). **d.** Mean ± SEM of the ΔPD between VIS and ROT blocks across all participants. * *P* < 0.05.

**Figure 5:**
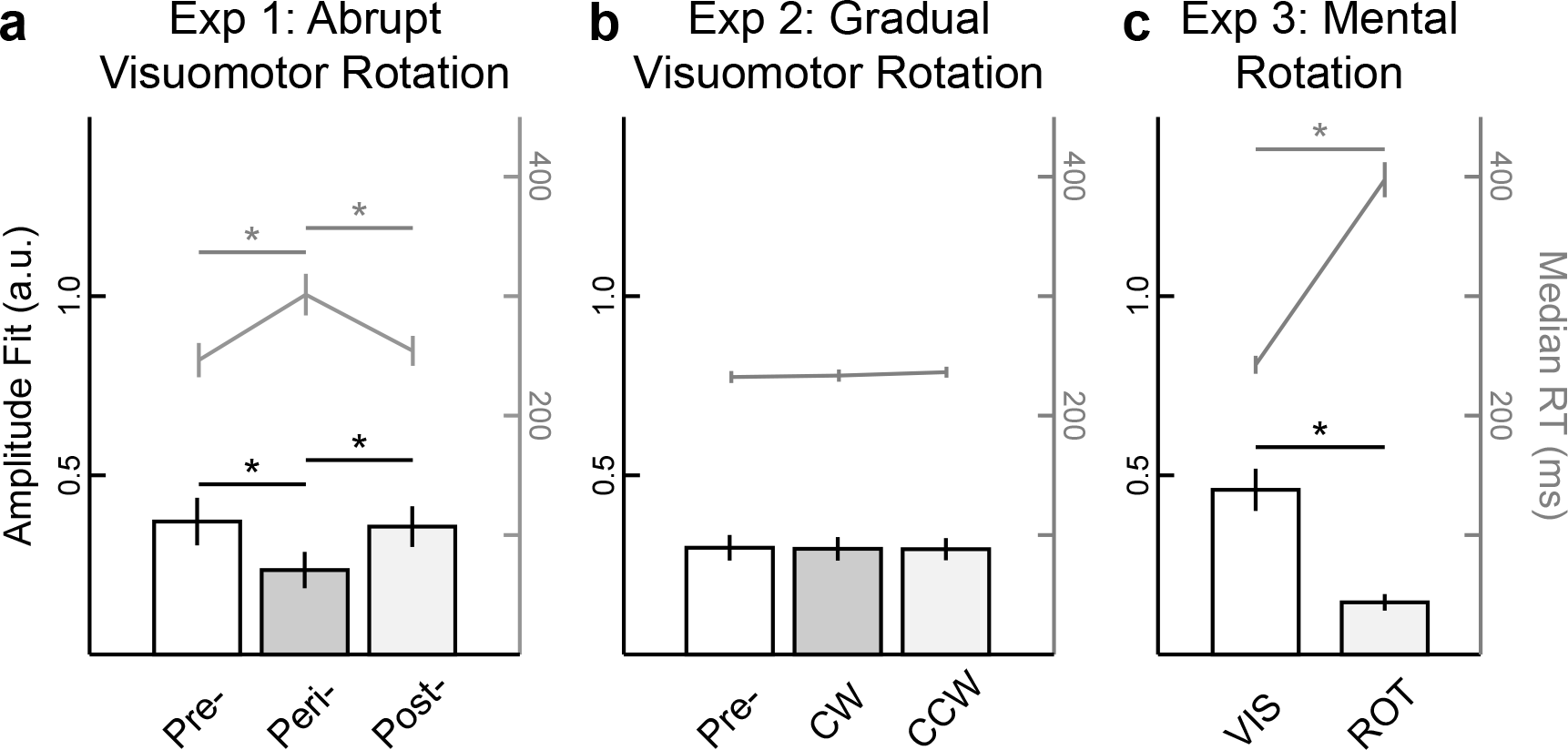
An explicit aiming strategy attenuated SLR magnitude and increased RTs a-c. Group mean ± SEM of both the amplitude parameter for the sinusoidal fits during the SLR epoch (bars, left axis) and median RTs (lines, right axis) across the three different experiments. * *P* < 0.05.

**Figure 4a** shows the endpoint reach direction from a participant who performed the ROT block first. There was no aftereffect during the initial few cycles after the end of the ROT block, which is consistent with the absence of implicit motor learning. **Figure 4b** shows a participant’s mean movement trajectories and PEC EMG activity for leftward and rightward stimulus locations (180° and 0° location, filled and open lines, respectively). Note that regardless of the voluntary movement direction, we observed greater EMG activity after leftward compared to rightward stimulus presentation during the SLR epoch in both the VIS (**Fig. 4b**, black lines, 2way ANOVA – direction and block, interaction effect, *F*_(1,225)_ = 12.57, *P* = 0.0005, post-hoc Tukey’s HSD, *P* < 10^−8^) and ROT blocks (red lines, post-hoc Tukey’s HSD, *P* < 10^−7^). Like the previous two experiments, we derived the PD of EMG activity during both the SLR and MOV epochs (**Fig. 4c**).

Across our sample, we observed a reliable shift in PD between the VIS and ROT blocks during the MOV epoch (**Fig. 4d**, ΔPD = 93.6° ± 1.5° CW, one sample *t*-test, *t*_12_ = 63.0, *P* < 10^−15^). In contrast, the SLR tuning did not reliably differ between the two blocks (ΔPD = −2.5° ± 3.8° CCW, one sample *t*-test, *t*_12_ = −0.7, *P* = 0.52). Although there was a significant attenuation in the amplitude of the SLR tuning curve between the VIS and ROT blocks (paired *t*-test, *t*_12_ = 5.96, *P* < 10^−4^), this attenuation was most likely due to the corresponding increase in RT during the ROT block, as SLR magnitude is known to decrease when preceding movements with longer RTs [15,20]. This decrease in amplitude was also observed during the Peri-Rotation block in Experiment 1, when there was also an increase in median RTs, but a decrease in amplitude was not seen in Experiment 2, when there was no reliable increase in median RTs (see **Fig. 5** for the relationship between SLR amplitude fits and median RTs in all three experiments). Thus, in Experiment 3, learning induced during a mental visuomotor task did not systematically alter the tuning of the SLR.

## DISCUSSION

Recent studies have suggested that motor learning can be driven by multiple learning components: an implicit learning component related to the mismatch between the actual and predicted sensory consequences of a generated motor command [5,9], and an explicit learning component that involves changes to aiming strategy [6,7]. What has not been clear from this literature is how such components engage various descending motor pathways. Here, we measured the changes in the directional tuning of EMG activity on the human pectoralis muscle during three variations of the visuomotor rotation task. We found both the implicit and explicit components of motor learning modulated the tuning of voluntary reach-related EMG activity. In contrast, we found that only the implicit motor learning component modulated the tuning of the earliest wave of muscle activity that is time-locked to the onset of a peripheral visual stimulus.

### Implicit motor learning drives the partial adaptation of SLR tuning during visuomotor rotations

Our central result is that implicit motor learning altered the directional tuning during the SLR epoch (85-125 ms after stimulus onset), while both implicit and explicit motor learning altered the tuning of reach-related MOV activity (−20 to 20 ms around RT, ~200-300 ms after stimulus onset). Thus, implicit motor learning can induce adaptation in the fastest, essentially reflexive, visuomotor pathway. The amount of adaptation was considerably less than either of our imposed visuomotor rotations: SLR tuning changed by 16.7° ± 3.6° for a 60° visuomotor rotation in Experiment 1, and by 12.8° ± 1.9° for an overall 40° visuomotor rotation in Experiment 2. These observations match well with previous indirect behavioral estimates of implicit learning component of ~10°-15° regardless of the magnitude of the imposed visuomotor rotation [6,21]. Such estimates are based on a subtraction logic, wherein the implicit component is estimated as the difference between the actual reach direction and the verbal reporting of the participant’s aiming direction.

The gradual visuomotor rotation used in Experiment 2 attempted to minimize the explicit aiming component of motor learning. Evidence that participants learned the new visuomotor mapping without using an explicit aiming strategy is found in the lack of difference in RTs between the veridical and rotation blocks (**Fig. 5**), and post-experiment confirmation that our participants were unaware of any changes in the visuomotor mapping during the experiment [29,30]. However, a previous study has reported impaired learning rates during a similar gradual visuomotor task when participants concurrently performed a cognitively demanding task [30], suggesting a distinction between explicit awareness and contribution of other forms of learning. This may explain why we only observed a partial adaptation of SLR tuning (~13°) compared to a full adaptation during the MOV epoch (~40°). Our paradigm was designed to test the influence of error-based learning, but may have also engaged reinforcement-based learning [32] as participants gauged their success in hitting the target. Indeed, reinforcement-based learning was likely engaged in all three Experiments. Previous studies have shown that changes in sensorimotor mapping can be driven purely by reinforcement learning [33,34], which can occur without awareness [35]. However, unlike implicit motor learning, reinforcement learning does not produce aftereffects [36], and as shown in Experiment 3, does not change SLR tuning.

### Distinct neural substrates for the implicit and explicit components of motor learning

To our knowledge, no previous animal neurophysiological or human imaging studies have described a neural correlate for partial adaptation during either a gradual or an abrupt visuomotor rotation task. Previous fMRI studies have shown that BOLD activity within the posterior parietal cortex (PPC) faithfully encodes visual stimulus location during the visuomotor rotation task, regardless of the ensuing reach direction [37,38]. Similarly, during saccadic adaptation, neurons within the lateral intraparietal cortex also encode visual stimulus location rather than saccadic endpoint [39]. Conversely, both fMRI and neurophysiological studies have shown that both premotor and primary motor cortices encode the final movement direction, regardless of the visual stimulus location [38,40–43]. Thus, the pattern of the modulation of SLR tuning is distinct from signals observed in either the PPC or motor cortices, which would presumably be relayed via corticospinal projections.

Previous clinical studies suggest that implicit and explicit components of motor learning have distinct underlying neural substrates. For example, even though patients with prefrontal lesions lacked any explicit awareness of changes during an abrupt visuomotor rotation task, they still partially adapted their reaching movements [10,11]. This result suggested that while the explicit aiming component is impaired, the implicit motor learning component is spared in such patients. Conversely, patients with cerebellar damage show impairment when adapting to novel environments [44–46], regardless of the size or how the perturbation is imposed [47,48]. While these patients can still compensate for the sensorimotor perturbations through either reinforcement learning [33,36] or the use of an explicit aiming strategy [8], they still had impaired implicit error-based learning [8,9,36] and displayed much smaller aftereffects after motor learning [49].

### A cerebellar influence on the tectoreticulospinal pathway

Given that the cerebellum has been strongly implicated in implicit motor learning, we surmise that the changes in SLR tuning observed in Experiments 1 and 2 are modulated via the cerebellum. How then could the cerebellum be altering this visuomotor mapping? We have speculated that the SLR is mediated by a tectoreticulospinal pathway [15,18,20], and there is substantial evidence for interaction between the cerebellum and the reticular formation. Consistent with cerebellar projections to the reticular formation [50–52], electrical stimulation to both human [53] and non-human primate [54,55] cerebellum evokes short-latency EMG response on upper limb muscles. These responses are still intact even after the inactivation of the contralateral primary motor cortex [55]. Further, the cerebellum receives an internal copy of the descending reticulospinal command from propriospinal neurons via the lateral reticular nucleus [56].

The (tecto)-reticulospinal pathway has also been implicated in other rapid motor responses such as the startReact effect [57–60], forced-RT paradigms [25,61], or corrective reach movements [62–64]. Our results, which demonstrate a selective influence of implicit motor learning on this descending pathway, may also explain the adaptation of these responses during various motor learning paradigms. For example, both startReact and corrective reach movements are modulated during motor learning induced by a force field [65,66] or, as studied here, a visuomotor rotation [67,68]. However, the contribution of implicit versus explicit components of motor learning was not considered in these paradigms. Here, by isolating EMG activity attributable to the tectoreticulospinal pathway and segregating the implicit and explicit components of motor learning, we can directly quantify the influence of different components of motor learning via the changes in the tuning of the SLR. Such an approach may be particularly useful for future work on motor learning in animal models to directly quantify implicit motor learning, serving as a benchmark for comparison with simultaneously recorded neural activity.

## Acknowledgments

This work was supported by operating grants from the Natural Sciences and Engineering Research Council of Canada (NSERC) to BDC [RGPIN-311680], PLG [RGPIN-238338], JAP [RGPIN-2015-06714], from the Canadian Institutes of Health Research to BDC [MOP-93796], a NSERC Canada Graduate Doctoral Scholarship to CG, and a salary award from the Canada Research Chairs program to JAP.

## Author Contributions

Conceptualization - CG, JAP, and BDC; Methodology - CG and PLG; Investigation - CG; Writing, Original Draft - CG and BDC; Writing, Review and Editing - JAP and PLG; Funding Acquisition - BDC; Resources - PLG; Supervision - JAP, PLG, and BDC;

## STAR METHODS

### Key Resources Table

| REAGENT or RESOURCE | SOURCE | IDENTIFIER |
| --- | --- | --- |
| Experimental Models: Organisms/Strains |  |  |
| Healthy human participants | University of Western Ontario | N/A |
| Software and Algorithms |  |  |
| Matlab | Mathworks | https://www.mathworks.com/ |

### Contact for Reagent and Resource Sharing

All requests for further information and resources should be directed to and will be fulfilled by the Lead Contact, Dr. Brian D. Corneil.

### Experiment Model and Subject Details

In total, we had 32 participants (21 males and 11 females, mean ± SD age: 25 ± 5 years old) perform at least one of the three experiments. All participants were self-declared right-handed expect for one left-handed male and four left-handed females, had normal or corrected-to-normal vision, and reported no current visual, neurological, and/or musculoskeletal disorders. Participants provided written consent, were paid for their participation, and were free to withdraw from any experiment at any time. All procedures were approved by the Health Science Research Ethics Board at the University of Western Ontario.

### Method Details

The apparatus, electromyographic (EMG) recording setup, and parts of the data analyses has been previously described [17,18,20].

#### Apparatus and kinematic acquisition

Briefly, in all three experiments, participants sat at a desk with their right elbow supported by a custom-built air-sled. They performed right-handed horizontal planar reaches while holding the handle of a planar robotic manipulandum (InMotion Technologies, Watertown, MA, USA). The *x*- and *y*-positions of the manipulandum were sampled and recorded at 600 Hz. A constant rightward load force of 5 N was applied throughout Experiments 2 and 3. No load was applied in Experiment 1. All visual stimuli were presented onto an upward-facing horizontal mirror, located just below the participant’s chin level, which reflected the display of a downward-facing LCD monitor with a refresh rate of 75 Hz. The precise timing of the peripheral visual stimulus onset on the LCD screen was determined by a photodiode. The mirror occluded view of the participant’s right arm throughout the experiment and real-time visual feedback of the handle of the manipulandum was given by a small red cursor on a white background.

#### EMG acquisition

EMG activity from the clavicular head of the right pectoralis major (PEC) muscle was recorded using either intramuscular (Experiment 1) or surface recordings (Experiment 2 and 3). Intramuscular EMG activity was recorded using fine-wire (A-M Systems, Sequim, WA, USA) electrodes inserted into the PEC muscle (see Wood et al., 2015 for insertion procedure). Briefly, for each recording we inserted two monopolar electrodes ~2.5 cm into the belly the PEC muscle. Insertions were aimed ~1 cm inferior to the inflection point of the clavicle, and staggered by 1 cm along the muscle’s fiber direction. All intramuscular EMG activity was recorded with a Myopac Junior System (Run Technologies, Mission Viejo, CA, USA). Surface recordings were made with doubled-differential electrodes (Delsys Inc., Natick, MA, USA) placed at the same location as the intramuscular recordings. EMG activity and the photodiode signal were digitized and recorded at 4 kHz.

#### Experiment 1: Abrupt visuomotor rotation task

Each trial began with the appearance of a central start position. Participants (*N* = 7/8 with a detectable SLR, SLR+, see below detection criterion) moved the cursor into the start position and after a randomized delay in the start position (1-1.25 sec) a peripheral black circle appeared (10 cm away from the start position at one of eight equidistant locations). The onset of the peripheral visual stimulus coincided with the offset of the start position. Participants were instructed to perform an out-and-back reach movement towards the peripheral stimulus. Additionally, they were instructed to reach as accurately as possible with the cursor to the peripheral stimulus during the outward movement. A small yellow circle also appeared at the position where the cursor crossed the 10-cm radius of the start position; this provided additional visual feedback on the accuracy of the outward reach movement.

Each participant performed 11 sub-blocks during the experiment, each sub-block consisted of 20 cycles (**Fig. 2a**, one cycle consists of eight trials, one trial for each of the eight different stimulus locations). In the first three sub-blocks (Pre-Rotation Block, black shade), the cursor veridically represented handle position. During the next four sub-blocks (Peri-Rotation Block, red), the cursor representing handle position was rotated by 60° CW around the start position. In the final four sub-blocks (Post-Rotation Block, blue) the cursor once again represented handle position.

#### Experiment 2: Gradual visuomotor rotation task

Like in Experiment 1, participants (*N* = 14/14 SLR+) moved the cursor into the start position and after a randomized delay in the start position (1-1.25 sec) a peripheral black circle appeared at one of eight equidistant locations around the start position. Participants were instructed to perform an out-and-back reach movement towards the peripheral stimulus and reach as accurately as possible with the cursor to the peripheral stimulus during the outward movement. However, during this task no yellow circle was presented after each outward reach movement.

Each participant performed nine sub-blocks, each consisting of 20 cycles (**Fig 3a**). In the first two sub-blocks (Test Block 1), the cursor veridically represented handle position. A gradual rotation was imposed during the third sub-block, in which the cursor representing handle position was rotated by 1° around the start position after each cycle; over the entire block the total rotation was 19°. During Test Block 2 (sub-blocks 4 and 5), participants performed reaches while the cursor was constantly rotated by 20°. In the next two sub-blocks (sub-blocks 6 and7), a gradual rotation was imposed 1° per cycle in the opposite direction as in sub-block 3; thus, by the end of sub-block 7 the total rotation imposed during the two sub-blocks was 39°. During Test Block 3 (sub-blocks 8 and 9), participants reached with a constant 20° rotation, which was in the opposite direction as Test Block 2. Participants were counterbalanced between experiencing either a CW or CCW rotation first (*N* = 7 per group, solid or dashed lines in **Fig. 3a**, respectively). Thus, all participants performed visually-guided reaches with veridical feedback (Pre-Rotation), and reaches with both a 20° CW and 20° CCW rotations (black, red, and blue shades in **Fig. 3a**, respectively).

#### Experiment 3: Mental visuomotor rotation task

Each trial began with the appearance of a start position and black outlines of the of eight equidistant locations 10 cm from the start position. Participants (*N* = 13/18 SLR+) moved the cursor into the start position and after a randomized delay in the start position (1-1.25 sec) one of the peripheral stimulus location was filled. Each participant performed six sub-blocks of 20 cycles (**Fig. 4a**). In three of the sub-blocks (VIS Block), participants performed out-and-back reach movements to the peripheral stimulus, while in the other three rotation sub-blocks (ROT Block), participants were instructed to reach towards the open stimulus location 90° CCW to the filled in peripheral stimulus location. Unlike Experiments 1 and 2, the cursor always veridically represented handle position throughout the experiment. The order of the blocks was counterbalanced between participants (*N* = 9 per group).

### Quantification and Statistical Analyses

#### Data pre-processing

All analyses were performed with custom-written scripts in Matlab (version R2014b, Mathworks Inc., Natick, MA, USA). To achieve sample matching between the kinematics and EMG data, all kinematic data was up-sampled from 600 Hz to 1000 Hz with a low-pass interpolation algorithm, and then lowpass-filtered with a second-order Butterworth filter with a cutoff at 150 Hz. Reach reaction times (RTs) were calculated as the time from the onset of the peripheral visual stimulus (measured by the photodiode) to the initiation of the reach movement. Reach initiation was identified by first finding the peak tangential movement velocity after stimulus onset, and then moving backwards to the closest time at which the tangential velocity profile surpassed 8% of the peak velocity. All EMG data was rectified and then either bin-integrated into 1 ms bins (intramuscular) or down-sampled (surface) to 1000 Hz. EMG activity was then normalized relative to each block’s mean baseline EMG activity (defined as the mean EMG activity 40 ms prior to the onset of the peripheral visual stimulus). We defined the SLR epoch as 85-125 ms after stimulus onset and the SLR magnitude as the mean EMG activity during the SLR epoch. We also defined the reach-related movement (MOV) epoch as 20 ms before to 20 ms after reach RT. All trials with RTs less than 185 ms were excluded to prevent contamination of the SLR epoch by shorter latency reach-related responses [18,20].

To determine the normalized movement trajectories, we first determined the movement duration for each trial individually. The movement duration was defined as the time when the handle position surpassed 2 cm from the center of the start position to 50 ms after the time when the handle position surpassed 8 cm from the center of the start position. We then interpolated the movement duration into 101 equally spaced time-samples, and calculated the *x*- and *y*-positions at each given time-sample.

#### SLR Detection and Latency Analysis

Based on previous studies detecting the presence of the SLR [15,69], we also used a receiver-operating characteristic (ROC) analysis to quantitatively detect the presence of a SLR. In all experiments, we examined EMG activity for leftward and rightward reaches during veridical visual feedback, and we performed the following ROC analysis. For every time-sample (1 ms bin) between 100 ms before to 300 ms after visual stimulus onset, we calculated the area under the ROC curve between the leftward and rightward trials. This metric indicates the probability that an ideal observer could discriminate the side of the stimulus location based solely on EMG activity. A ROC value of 0.5 indicates chance discrimination, whereas a value of 1 or 0 indicates perfectly correct or incorrect discrimination, respectively. We set the thresholds for discrimination at 0.6; these criteria exceed the 95% confidence intervals of data randomly shuffled with a bootstrap procedure [70]. The earliest discrimination time was defined as the time after stimulus onset at which the ROC was above 0.6 and remained above that threshold for at least 5 out of the next 10 samples. Previous studies have also reported decreased SLR magnitude during an anti-reach task [20], thus we lower our threshold to 0.55 for the ROT block in Experiment 3. Based on the ROC analyses we defined the SLR epoch as from 85 to 125 ms after visual stimulus onset and categorized any participant with a discrimination time <125 ms as having a SLR (SLR+ participant). Across the three experiments we could reliably detect a SLR in 29 out of 32 participants.

#### Tuning curve fit

To determine the tuning curve of EMG activity during both the SLR and MOV epochs, we assumed that the relationship between EMG activity and the peripheral visual stimulus location took the form of a sinusoidal function Eq. 1:

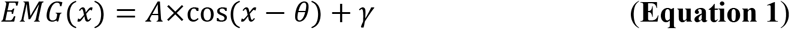

in which *x* is the angular location of the peripheral visual stimulus in degrees; *EMG(x)* is the logarithm of the normalized EMG activity for the given stimulus location; *A* is the amplitude of the sinusoidal fit; *θ* is the preferred direction (PD) of the sinusoidal fit; and *γ* is the offset of the sinusoidal fit. We used Matlab’s curve fitting toolbox, in which we constricted our parameters so that *A* < 0 and 0 ≤ *θ* < 360, and the starting point of the parameters were *A* = 1, *θ* = 180°, and *γ* = 0.

#### Statistical Analyses

All statistical analyses were performed using either a paired *t*-test or repeated-measure ANOVA. For all post-hoc, we used a Tukey’s HSD correction. The statistical significance was set as *P* < 0.05.

### Data and Software Availability

All data was analyzed using MATLAB R2014b.

